# Multiplexing of functionally distinct inputs by intrinsic modulation of spike timing in a monoaminergic nucleus

**DOI:** 10.1101/2024.09.09.611991

**Authors:** Michael Lynn, Leonard Maler, Jean-Claude Béïque

**Affiliations:** Department of Cellular & Molecular Medicine, University of Ottawa

## Abstract

Monoaminergic nuclei such as the serotonergic dorsal raphe nucleus (DRN) receive synaptic inputs containing functionally distinct streams of information, yet the dimensionality of the resulting output population code and its cellular underpinning are currently unknown. By combining electrophysiological and computational approaches, here we uncover separable neural encoding of two excitatory inputs conveying disjunct information to DRN 5-HT neurons - the lateral habenula (LHb) and medial prefrontal cortex (mPFC). Dual-color opsin strategies revealed that a population of 5-HT neurons receive inputs from both mPFC and LHb. Subthreshold excitatory postsynaptic potentials triggered by both inputs were largely indistinguishable, yet suprathreshold spiking behavior exhibited input-specific latencies and dispersion statistics. A support vector machine classifier demonstrated that input identity can be accurately decoded from spike timing, but not subthreshold events, of under ten 5-HT neurons. Upon examining the intrinsic cellular mechanisms in 5-HT neurons that couple EPSPs to spiking dynamics, we uncovered two likely candidate mechanisms: a low-threshold calcium conductance that selectively boosts slow excitatory inputs, and a subthreshold, voltage-dependent membrane noise that generates variation of spike latency and jitter. Stochastic simulations suggest that these two intrinsic properties of 5-HT neurons are sufficient to transform LHb and mPFC inputs into distinct output spiking patterns. These results reveal that hub-like networks like the DRN can segregate distinct informational streams by a cell-intrinsic mechanism. The resulting emergent population spike synchrony code provides a means for the DRN to widely broadcast these streams as a multiplexed signal.

**Significance statement:** Phylogenetically old neuromodulatory systems in the brain, such as the serotonergic dorsal raphe nucleus, are compact yet richly innervated structures. Here, we use the raphe as a testbed to ask how distinct informational sources to hub-like networks are processed or integrated into a coherent neural code. Using electrophysiological and computational methods, including biophysically grounded stochastic simulations, we find that intrinsic noise mechanisms in serotonergic neurons are critical to transform approximately matched subthreshold excitation into distinct spike timing profiles. Thus, cell-intrinsic noise mechanisms can effectively synthesize a spike synchrony code that, we hypothesize, multiplexes input information to hub-like networks at the population level even in the absence of strong local circuit interactions.

## Introduction

The phylogenetically old monoaminergic nuclei share common core organizational features. They are anatomically compact, give rise to widespread and distributed axonal projections, and are controlled by expansive and convergent sets of synaptic inputs carrying distinct informational content. A corollary computational question is how these multiple, temporally overlapping, competing inputs are differentially processed by these hub networks. In principle, populations of neurons in these networks could either collapse independent streams of information into a low-dimensional rate code, or alternatively could independently encode features of the inputs in a way that can be decoded by downstream regions, a neural strategy termed ‘multiplexing’ which includes a diversity of rate or temporal code implementations (Naud & Sprekeler, 2018; Payeur et al., 2021; Oswald et al., 2004; Clarke & Maler, 2017; Lankarany et al, 2019; Ratté et al., 2013; Friedrich et al, 2004; Stringer et al., 2019; Nogueira et al., 2023). Physiological mechanisms for multiplexing can involve cell-intrinsic properties of individual neurons (Doiron et al., 2002; Laing et al., 2003), or alternatively can depend on large-scale network interactions, generating multiplexed neural codes at the population level (e,g, Stringer et al., 2019). It is currently unknown whether monoaminergic nuclei functionally discriminate competing excitatory inputs, an obligatory prerequisite of multiplexing, as well as what potential neural coding strategies could implement such a discrimination.

The midbrain dorsal raphe nucleus (DRN) is a monoaminergic nucleus that has received sustained attention notably because of its role in mood regulation and decision-making processes (Warden et al., 2012; Ren et al,. 2018; Li et al, 2016; Paquelet et al., 2022; Cohen et al., 2015). The anatomical and functional organization of DRN is complex, with DRN 5-HT neurons innervating a vast array of cortical and subcortical structures and receiving broad inputs, including from structures encoding cognitive and valence information such as the medial prefrontal cortex (mPFC) and lateral habenula (LHb) (Dorocic et al., 2014; Weissbourd et al., 2014; Geddes et al., 2016; Zhou et al., 2017). These two inputs provide excitatory synaptic inputs to partially overlapping anatomical divisions (Zhou et al., 2017) including the ventral section of DRN, yet are functionally distinct, with mPFC representing persistent cognitive states (Miller & Cohen, 2001; Bari et al., 2019) and LHb cells responding to transient negative reward prediction errors with high temporal precision spiking responses (Matsumoto & Hikosaka, 2007; Bromberg-Martin & Hikosaka, 2011).

Because of their fundamentally distinct informational content, we reasoned that the inputs from the PFC and the LHb present an ideal testbed to explore discriminative neural strategies in the DRN. We thus compared the LHb and PFC inputs to DRN 5-HT neurons using cellular electrophysiology and optogenetic approaches. While individual 5-HT neurons received broadly similar excitatory synaptic inputs from both sources, those from the PFC evoked spikes in 5-HT neurons with distinct statistics (*i.e.,* latency and jitter) than those triggered by LHb input activation. Machine learning approaches demonstrated that these timing differences were theoretically sufficient for downstream circuits receiving projections from DRN populations to functionally discriminate these input streams, suggesting a multiplexed population code for LHb and PFC input information. Physiological investigations and stochastic simulations defined a cell-intrinsic mechanism involving a low-threshold calcium channel-mediated conductance in 5-HT neurons that is responsible for generating these spike timing differences. Taken together, these data suggest that the phylogenetically old, expansively projecting DRN 5-HT system can broadcast multiple streams of information that are reflecting the contribution of distinct source inputs.

## Results

### Single 5-HT neurons integrate multiple long-range inputs

We first sought to determine whether LHb and mPFC inputs synapse onto distinct, non-overlapping populations of 5-HT DRN neurons or, alternatively, whether some subpopulations receive common input. To probe the integration of mPFC and LHb inputs at single 5-HT neurons, we employed a dual-color opsin approach (Klapoetke et al., 2014) wherein we bilaterally injected adeno-associated viruses (AAVs) expressing Chronos (excited by blue light) in mPFC and Chrimson (excited by red light) in LHb (Fig. 1A). Using an *in vitro* slice physiology preparation, we recorded in whole-cell configuration from genetically identified 5-HT neurons (SERT-CRE-tdTomato mouse line; see Methods) primarily in the ventromedial and dorsomedial regions of the DRN. In voltage clamp mode (Vm=-70 mV), brief (1 ms) pulses of blue (473 nm) or red (550 nm) light were delivered to the slice through an optic fiber. In agreement with previous work (Geddes et al., 2016; Zhou et al., 2017), we observed reliable inward photocurrents (excitatory post-synaptic potentials, EPSCs) which were time-locked to optical stimulation (Fig. 1B), indicating the presence of long-range glutamatergic synaptic inputs from LHb or mPFC. Single 5-HT neurons showed a diversity of responses, responding to neither, one, or both wavelengths of light. Overall, 43.8% of 5-HT neurons responded to both wavelengths of light, 18.8% of neurons responded to neither, and the remainder responded to only one wavelength (Fig. 1C).

**Figure 1:**
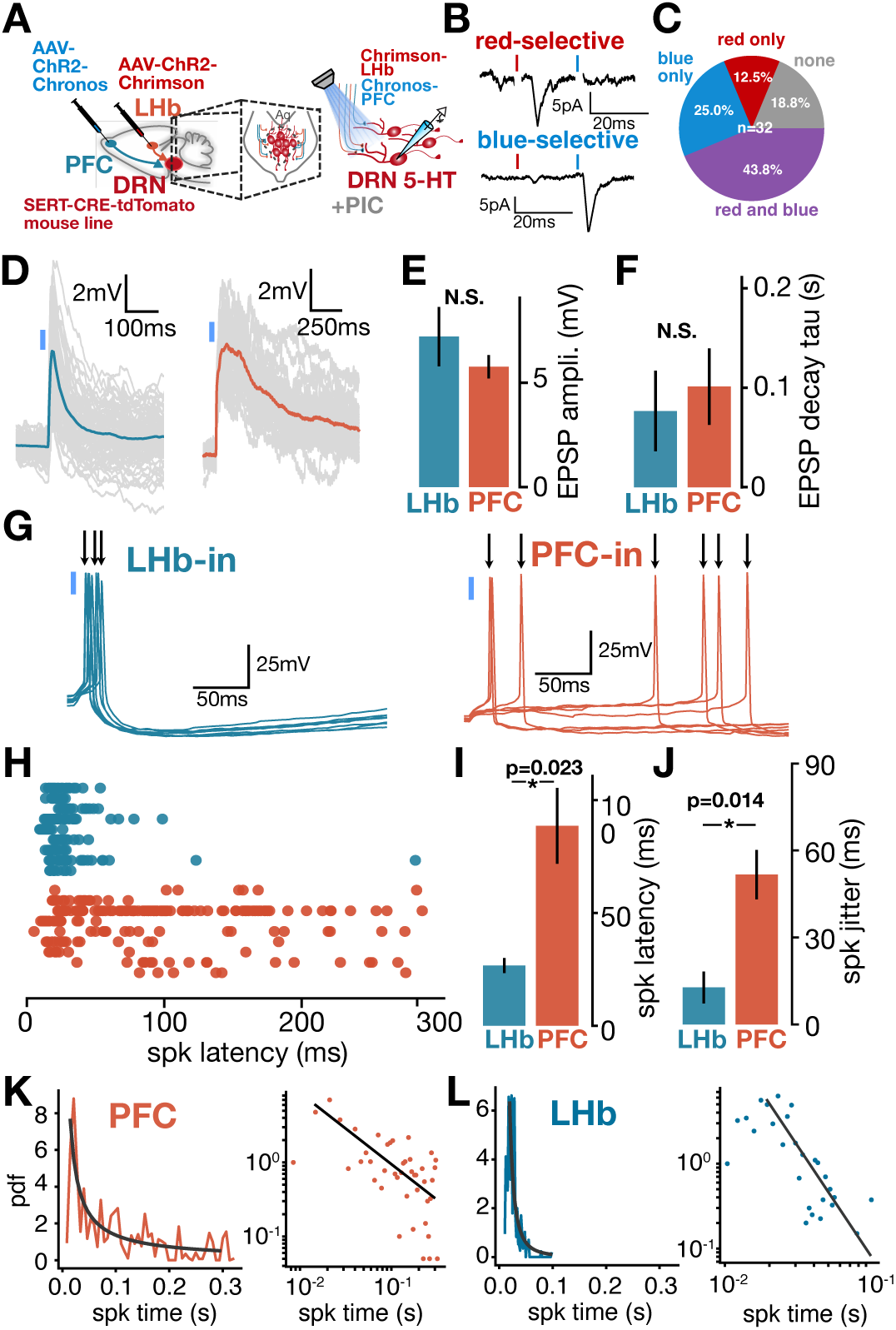
Subthreshold and spike timing characteristics of long-range PFC and LHb input to 5-HT neurons. **A**: Experimental setup showing dual-color opsin strategy. **B**: Example voltage-clamp traces of individual 5-HT neurons responding only to red (top) and blue (bottom) photostimulation. **C**: Quantification of fraction of neurons responding to each light type (n=32). **D**: Example current-clamp traces of EPSPs of individual 5-HT neurons in response to photostimulation of PFC (red) or LHb (blue) inputs. **E-F**: Amplitude and decay (mean +-SE) of EPSPs in individual neurons in response to PFC (red, n=9 neurons) or LHb (blue, n=9 neurons) input (paired T-test). **G**: Example current-clamp traces of spiking responses of individual 5-HT neurons in response to LHb (blue) or PFC (red) photostimulation. **H**: Spike timing of individual neurons (rows) in response to LHb (blue, n=9 neurons) or PFC (red, n=9 neurons) photostimulation. **I-J**: Spike latency and jitter (mean +-SE) for individual neurons in response to LHb (blue, n=9 neurons) or PFC (red, n=9 neurons) photostimulation (paired T-test). **K**: Probability distribution of spike times in response to PFC photostimulation across all neurons (left) and plotted as a log-log relationship with line of best for power-law scaling (right). **L**: Probability distribution of spike times in response to LHb photostimulation across all neurons (left) and plotted as a log-log relationship with line of best fit for power-law scaling (right).

To rule out overlap between the excitation spectra of these two opsins, we delivered a set of patterned dual-color stimuli talking advantage of opsin desensitization, similar to Christoffel et al (2021) (see Methods). Briefly, if the same population of synapses expressing a single opsin are activated twice by paired-pulse stimulation, this should lead to release probability changes and measurable short-term dynamics, either facilitation due to presynaptic calcium accumulation, or depression due to receptor desensitization or vesicle depletion (Fig. S1A, top; Dobrunz & Stevens, 1997). However, if sequential pulses each activate distinct synaptic populations (*e.g.* populations expressing two spectrally distinct opsins), then no short-term dynamics should be possible. Consistent with this idea, we delivered paired-pulse stimulation at 20 Hz either with two red pulses (activation of the same synaptic population expressing Chrimson), or with a sequential red and blue pulse (activation of two synaptic populations expressing either Chrimson or Chronos). The red-red paired-pulse stimuli triggered prominent short-term depression (Fig. S1A, top), while the red-blue paired-pulse stimuli triggered minimal short-term dynamics, as assessed through comparison with the event amplitude of a solitary blue photostimulation (Fig. S1A, bottom). Paired comparison across the population indicated that the same neurons exhibited minimal dynamics with red-blue photostimulation (paired-pulse ratio close to 1) compared to red-red photostimulation (paired-pulse ratio < 1; Fig. S1B) This indicates an effective separation of the excitation spectra of both opsins in this subset of dual-responder neurons. Single-opsin controls (Chrimson in LHb alone; Fig. S1C-D) revealed that while blue light could occasionally activate the red opsin, our patterned dual-color stimulation accurately discriminated this condition, with similar depression dynamics observed in red-red paired-pulse and red-blue paired-pulse stimulation as expected (compare Fig. S1D with Fig. S1B). Together, these results indicate that potential misidentification of input specificity due to spectral overlap of complementary opsin pairs can be effectively screened with appropriately designed stimuli. While this overall strategy does not lend itself with ease to precise quantitative estimates, it nonetheless shows that at least a subpopulation of DRN 5-HT neurons receive joint long-range excitatory input from both mPFC and LHb.

### Subthreshold responses of DRN 5-HT neurons to mPFC and LHb inputs are similar

In principle, the information content encoded by distinct mPFC and LHb inputs to individual neurons may be collapsed into a single low-dimensional code or, alternatively, information from each input may be preserved. To address this question, we first investigated synaptic features of PFC and LHb inputs to 5-HT neurons. We injected an AAV expressing channelrhodopsin (ChR2) bilaterally into either LHb or mPFC in two groups of mice, and conducted whole-cell recordings from identified 5-HT neurons (see Methods). Brief pulses of blue light triggered fast inward currents, in keeping with previous reports showing monosynaptic, glutamatergic connections between PFC and LHb axons and DRN 5-HT neurons (Fig. S2A-B; Vm=-70mV). In current-clamp mode, photostimulation triggered time-locked excitatory postsynaptic potentials (EPSPs) that were broadly similar between LHb and mPFC inputs, although those induced from PFC inputs often displayed longer decay (Fig. 1D). Indeed, while the mean amplitude and decay of EPSPs across cells were not statistically different between inputs (Fig. 1D-F), the distribution of EPSP amplitudes and decays across trials was slightly but significantly different between PFC and LHb evoked events (Fig. S2C-D). Together, these results indicate that 5-HT neurons receive long-range inputs from LHb and mPFC that are approximately matched in amplitude and timecourse at subthreshold membrane potentials.

### A spike-timing code discriminates input identity from mPFC and LHb in single 5-HT neurons

We next examined how the EPSPs from the PFC or the LHb are transformed into spiking output in individual 5-HT neurons. We optogenetically stimulated long-range inputs from either mPFC or LHb while recording from 5-HT neurons in current clamp, and examined in details the resulting sparse spiking responses elicited by each input separately. By fine-tuning direct current injection, we ensured: 1) comparable resting membrane potential (Fig. S2E), 2) comparable fraction of trials eliciting a spike (Fig. S2F) between neurons and; 3) that spontaneous spikes (*i.e.,* occurring before the optical stimulation) were not present. LHb stimulation triggered low-latency spiking with very little variability (jitter) between trials, largely consistent with theories of spike times encoding the derivative of inputs - in this case, EPSPs with sharp rise-times (Mainen & Sejnowski, 1995). mPFC stimulation, in contrast, triggered spikes with much longer latencies with high jitter (Fig. 1G-I). The latencies of the mPFC triggered spikes, often lasting hundreds of milliseconds, were qualitatively similar across all 5-HT neurons measured (Fig. 1H). In contrast, LHb-triggered spikes exhibited stereotyped, sharp voltage trajectories to spike threshold that were highly reliable across trials, while in contrast PFC-triggered spikes exhibited longer voltage trajectories that demonstrated less trial-to-trial reliability (p=0.014; Fig. S2G-J). mPFC inputs yielded spike-times that were distributed with a heavy right tail that could be well fit with a power-law decay (Fig. 1K), while LHb inputs triggered spike-times that could also be fit with a power-law decay, but with a larger negative exponent reflecting steeper decay (Fig. 1L). However, since spike-times spanned less than one order of magnitude we could not formally distinguish this power-law decay from an exponential decay (Fig. S2K-N). This is consistent with theories of fluctuation-driven activity regimes, where spike timing depends on membrane potential and noise statistics (Gerstein & Mandelbrot, 1964; van Rossum et al., 2003).

Could differential stimulus-driven spiking for each input reflect an effective segregation of each informational stream within populations of 5-HT neurons? One compatible theory is that of synchrony-division multiplexing (Lankarany et al., 2019; Ratté et al., 2013; Friedrich et al, 2004) suggesting that synchronous firing of neurons in a population can carry information distinct from the time-averaged firing rate of individual neurons. Our data suggests that the precisely time-locked response of many 5-HT DRN neurons to LHb inputs would trigger synchronous spiking across the population. In contrast, mPFC input is expected to generate asynchronous spiking of 5-HT neurons. These two firing patterns could, in principle, be decoded by circuits downstream of 5-HT neurons. To explore this possibility, we first applied linear classifiers to our spike timing data from each input, with the aim of determining whether the identity of the input (LHb or mPFC) could be theoretically determined solely from the spiking output of a population of 5-HT neurons. We generated synthetic training and test datasets (5-fold cross-validation, see Methods) based on an unbiased sampling of experimentally-derived spike-times from single trials for each input (Fig. 2A). By training a support vector machine classifier and assessing performance on held-out data, we quantified how spike timing-based decoding performance varied as a function of the number of spikes. Surprisingly, we found that spike timing information provided by as few as 6 neurons allowed good (>90%) decoding of the identity of the synaptic input driving these spikes (i.e., mPFC or LHb) (Fig. 2B-C), indicating that synchrony/asynchrony-based decoding could be biologically plausible even by areas receiving sparse innervation by 5-HT axons. We next determined whether the EPSPs alone (i.e., subthreshold voltage amplitudes or decays) from both inputs could be discriminated, and found poor decoding for both inputs, not exceeding 90% correct classification with >30 observations (Fig. 2D-G). The poor discriminability between the EPSPs triggered by PFC and LHb inputs onto 5-HT neurons stands in stark contrast to the robust discriminability of 5-HT neuron’s output (based on spike timing) induced by these same PFC and LHb inputs. Thus, a latent cellular feature operant in 5-HT neurons is likely responsible for transforming broadly indistinguishable synaptic inputs into spiking outputs with substantial timing differences.

**Figure 2:**
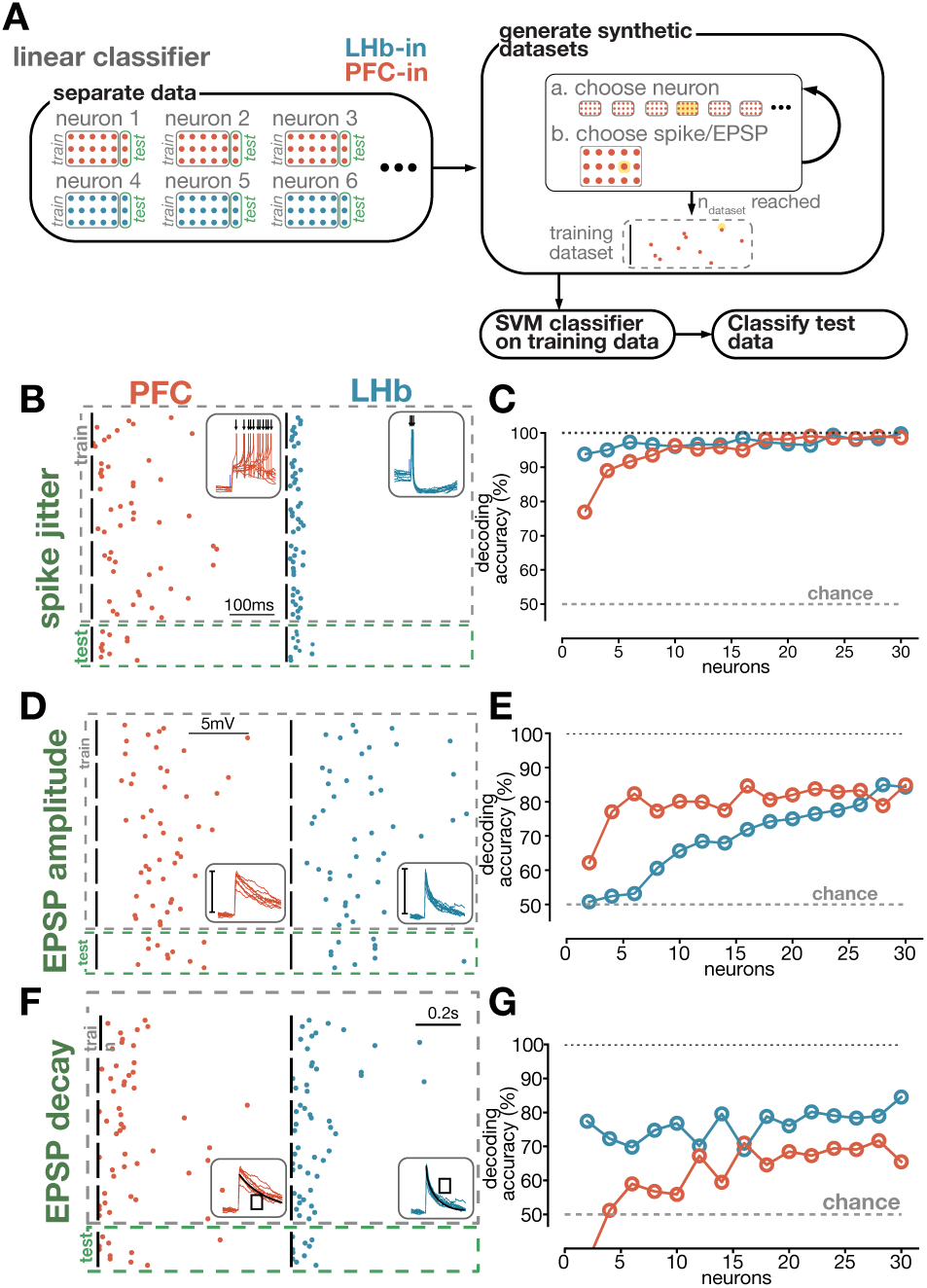
Decoding long-range inputs by spike-timing and subthreshold features in 5-HT populations. **A**: Schematic showing approach for generating synthetic datasets and applying a support vector machine linear classifier to data (5-fold cross-validation). **B**: Example synthetic datasets of spike times from PFC (red) or LHb (blue). Each vertical line represents one example dataset of n=10 spike times. **C**: Decoding accuracy of PFC (red) and LHb (blue) synthetic datasets from linear classifier as a function of dataset size (neurons). Dotted line indicates chance decoding performance (50%). **D**: Example synthetic datasets of EPSP amplitude from PFC (red) or LHb (blue). Each vertical line represents one synthetic dataset of n=10 EPSP amplitudes. **E**: Decoding accuracy of PFC (red) and LHb (blue) synthetic datasets from linear classifier as a function of dataset size (neurons). Dotted line indicates chance decoding performance (50%). **F**: Example synthetic datasets of EPSP decay from PFC (red) or LHb (blue). Each vertical line represents one synthetic dataset of n=10 EPSP amplitudes. **G**: Decoding accuracy of PFC (red) and LHb (blue) synthetic datasets from linear classifier as a function of dataset size (neurons). Dotted line indicates chance decoding performance (50%).

### A Calcium channel-mediated conductance expressed in 5-HT neurons support differential spike timing

We next sought to determine candidate cellular mechanisms in 5-HT neurons that could generate reliable differences in spike timing between inputs. It has previously been shown that low-threshold (T-type) calcium channels are highly expressed in several key subgroups of DRN 5-HT neurons (Templin et al., 2012; Okaty et al., 2020) and trigger rebound spiking in a subset of putative 5-HT neurons (Burlhis & Aghajanian, 1987). We first examined their expression in genetically identified 5-HT neurons and found that voltage steps to Vm = −40 mV triggered delayed inward currents lasting several hundred milliseconds, which were blocked by application of NiCl, a low-voltage calcium channel blocker (Fig. 3A; Lee et al., 1999, see Methods). The inward current was voltage-dependent, with a protracted rise-time and decay consistent with reported values for a low-threshold calcium channel (Fig. 3A-B; Perez-Reyes, 2003). Together, these data indicate that low-threshold calcium channels are expressed in 5-HT neurons and are activated at depolarized voltages.

**Figure 3:**
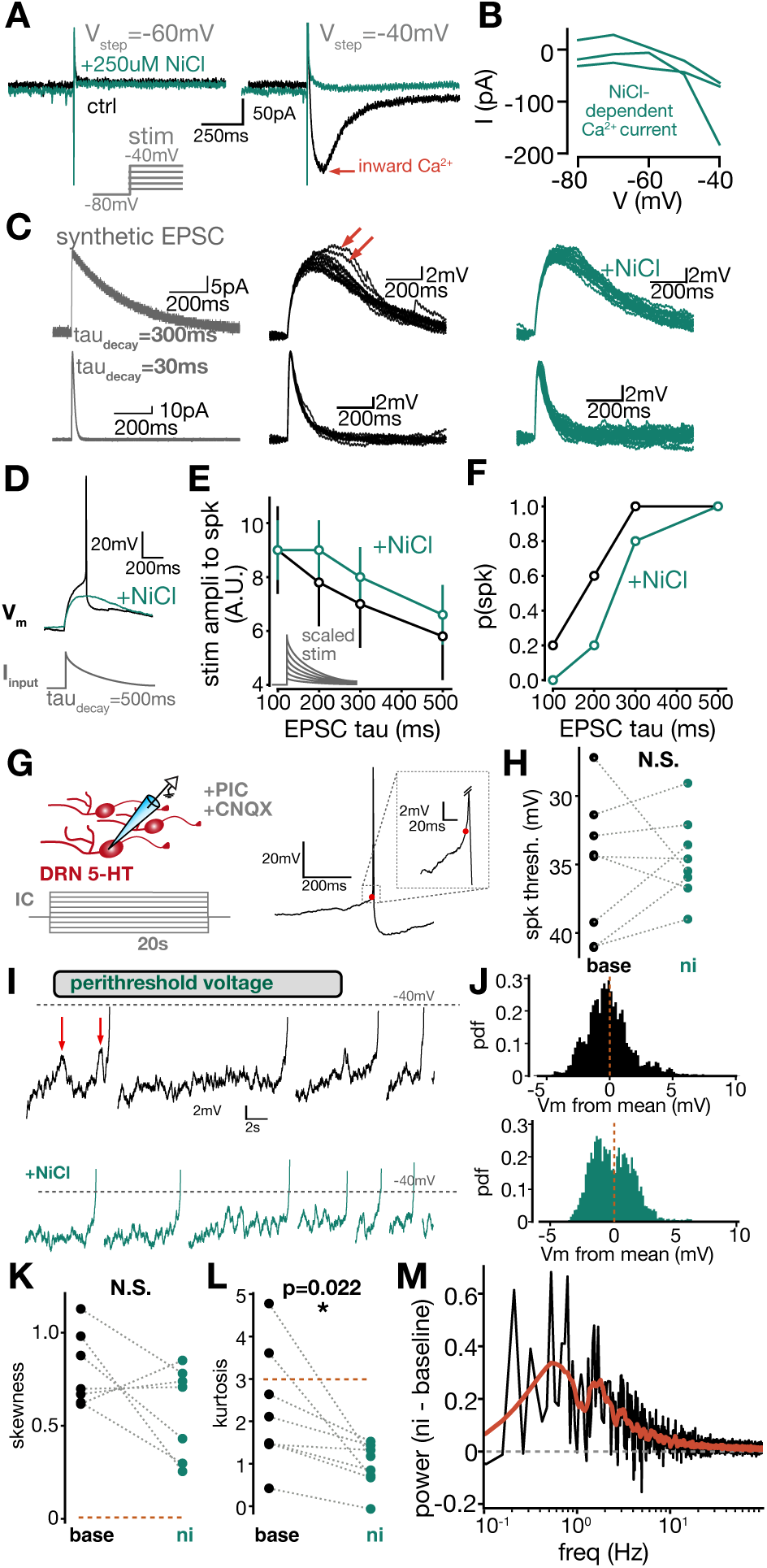
An inward calcium conductance in 5-HT neurons shapes EPSPs and contributes to membrane noise. **A**: Example traces in voltage-clamp showing an inward conductance at −40mV (right) but not −60mV (left) that is blocked by bath application of NiCl (blue trace). **B**: Voltage-dependence of the putative inward calcium conductance (n=3 neurons). **C**: Synthetic EPSCs injected in current-clamp into 5-HT neurons (left) produce transient voltage excursions (middle) blocked by bath application of NiCl (right). **D**: Example trace showing modulation of synthetic EPSC-triggered spiking by bath application of NiCl. **E**: Relationship between synthetic EPSC amplitude required to elicit a spike, and synthetic EPSC tau, in baseline conditions (black) and after bath application of NiCl (blue) (n=5 neurons, bars represent S.D.). **F**: Relationship between probability of obtaining an action potential across neurons, and synthetic EPSC tau, in baseline conditions (black) and after bath application of NiCl (blue) (n=5 neurons). **G**: Experimental setup for membrane noise experiments presented in **H-M** (left), and example of spike threshold estimation using fractional first-derivative method (right). **H**: Spike threshold is not modulated by bath application of NiCl (n=8 neurons, paired T-test). **I**: Example traces of a single neuron recorded in current-clamp during injection of constant perithreshold depolarizing current, before (top) and after (bottom) application of NiCl. Red arrows depict large voltage excursions (‘blips’). **J**: Probability distribution of membrane potentials for an example neuron before (top) and after (bottom) application of NiCl. **K**: Membrane potential distribution skewness is not significantly affected by bath application of NiCl (n=7, paired T-test). **L**: Membrane potential distribution kurtosis is significantly reduced by bath application of NiCl (n=8, paired T-test). **M**: Relative spectral power of calcium conductance, calculated by subtracting baseline membrane potential spectral power from post-NiCl washon membrane potential spectral power (n=8 neurons, mean shown.)

Next, we sought to examine how these low-threshold calcium channels modify incoming synaptic inputs of variable kinetics. To parametrize this effect, we injected in genetically identified 5-HT neurons synthetic EPSC waveforms which resembled incoming EPSCs with varying decay kinetics (Fig. 3C, left). The injection of synthetic EPSCs with slow decay kinetics led to variable voltage responses, both in terms of amplitude and decay, which we hypothesized may be due to stochastic and transient calcium channel activation (Fig. 3C, middle). Consistent with this hypothesis, bath application of low concentrations of NiCl selectively blocked the slow and high amplitude subset of voltage responses triggered by the slow synthetic EPSCs (Fig. 3C, right; also see Supp. Fig. S3). In contrast, charge-normalized synthetic EPSCs with faster decays trigered low variability voltage responses that lacked prolonged, high-amplitude events and that were not affected by application of NiCl (Fig. 3C; also see Supp. Fig. S3).

Since the activation of calcium channels and other ionic conductances can affect the timing of spikes (Cain & Snutch, 2013; van Rossum et al., 2003), we hypothesized that these channels might play a similar role in regulating spike timing in 5-HT neurons. Blocking ionic conductances should, in principle, increase the neuron’s membrane resistance and make it easier for a fixed magnitude of current input to depolarize the neuron sufficiently to trigger spikes. However, we found, surprisingly, that bath application of NiCl could reduce or even eliminate spiking response for a fixed perithreshold current input (Fig. 3D). This may indicate complex interactions between voltage events and activation/inactivation kinetics of this calcium conductance, as well as spatial profiles of channel expression throughout the dendrite and soma. We used synthetic EPSCs to explore how the production of spikes is tuned by calcium channels across the parameter space of event kinetics. The decay kinetics and amplitudes of the injected synthetic EPSCs were systematically varied while we recorded the voltage response of 5-HT neurons. We found, consistent with our earlier findings (Fig. 3D), that NiCl application increased the amplitude of the synthetic EPSC required to elicit spiking behavior, especially at long decay tau values (Fig. 3E) and reduced spiking probability across all neurons at the same fixed EPSC amplitude (Fig. 3F). These results highlight an important role of this calcium conductance in regulating the spiking output of 5-HT neurons.

We next sought a mechanistic explanation for how the activation of this peri-threshold conductance could generate variation in spike timing, thus plausibly modulating network synchrony. To this end, we adopted a dynamical approach and determined how voltage noise statistics – a determinant of spike timing-are modulated by these channels. We conducted experiments where finely spaced, long (20s) current steps were injected into identified 5-HT neurons before and after application of NiCl (Fig. 3G). Throughout these experiments, a cocktail of synaptic blockers was applied (see Methods). To focus on sub- and perithreshold regimes, spikes were identified and removed from the recordings using a fractional first-derivative method (Fig. 3G; see Methods; Trinh et al., 2019; Azouz and Gray, 2000). At perithreshold membrane potentials (defined by the first depolarizing current step eliciting any action potentials), we observed prominent voltage noise as well as depolarizing, non-spike-generating, events lasting ∼1s (Fig. 3I-J, top). We suspected that these slow depolarizing events, as well as fast voltage noise, were partially generated by stochastic activation and inactivation of these low-threshold calcium channels. Consistent with this, application of NiCl modulated the amplitude of both the slow events and fast voltage noise, and decreased the tail size of the membrane voltage distribution (Fig. 3I-J, bottom) without altering spike threshold per se (Fig. 3H).

To quantify these observations across the population of recorded 5-HT neurons, we measured the skewness (third central moment, indicating asymmetry) and kurtosis (fourth central moment, indicating tailedness) of the membrane potential distribution, before and after application of NiCl. Before NiCl application, the membrane voltage distribution had skewness values above 0, indicating a right tail, that was not affected by NiCl application (Fig. 3K). The distribution’s kurtosis was, however, significantly reduced by NiCl application, consistent with this calcium conductance being causally implicated in the generation of depolarized voltage excursions (Fig. 3L). To determine whether these effects were localized to perithreshold voltage regime, we considered the voltage dynamics during the first subthreshold current step (i.e., without spikes) and found no effect of NiCl bath application on kurtosis or skewness (Supp. Fig. S4). This indicates that the effects of low-threshold calcium channel activation on membrane dynamics is preponderant at perithreshold membrane potentials. Finally, we performed a fast Fourier transform of the time-varying perithreshold membrane potential (again with spikes removed) to determine how calcium channel blockade affected the frequency content of the noise. This analysis revealed a NiCl-dependent boost in frequencies in a 0.1-10Hz window (Fig. 3M). Collectively, these results indicate that low-threshold calcium channels robustly modulate membrane noise around spike threshold in 5-HT neurons.

We next naturally hypothesized that this perithreshold calcium conductance is causally involved in generating mPFC-evoked delayed spiking in 5-HT neurons. We thus repeated the photostimulation experiments of mPFC axons in the DRN and found that it triggered high-latency spikes (Fig. S5A, left), replicating our previous results (Fig. 1G). We then applied in these same recordings voltage steps to −45mV that triggered inward currents that were abolished by bath application of NiCl (Fig. S5B). We then resumed optogenetic stimulation of mPFC axons and found that the resultant spikes displayed noticeably shorter latency and less jitter (Fig. S5A, right; Fig. S5C). The effect of bath application of NiCl on reducing spike latency was statistically significant across the population of 5-HT neurons recorded (Fig. S5D-E, p=0.045, Kolmogorov-Smirnov test on spike times compiled across the population). Taken together, these results establish a causal relationship between the calcium channel conductance expressed in 5-HT neurons, and the delayed spike timing behavior evoked by long-range mPFC input.

### Dynamical regimes for synchronous and asynchronous spike generation

We next employed stochastic, biophysically grounded simulations to obtain a mechanistic understanding of how calcium channels could regulate the initiation time of action potentials, and thus modulate spike synchrony at the population level. We first considered a simplified scenario where Ornstein-Uhlenbeck noise (simulating barrages of synaptic input and/or intrinsic noise, see Methods; Destexhe et al., 2001) was injected to a toy neuron model with subthreshold membrane potential. This produced a voltage signal that evolved over time in a pseudo-random fashion (Fig. 4A). The time at which the voltage signal first crossed a depolarized spike threshold (called the first-passage crossing time; yellow dot in Fig. 4A) was recorded over 20,000 stochastic, independent simulations, generating a probability distribution of spike times (Fig. 4B). These simulations effectively correspond to a population of 5-HT neurons receiving a fixed synaptic input, and responding stochastically due to cell-intrinsic mechanisms. This approach allowed us to ask how the spike timing distributions we observed could be related to simple interactions between a stochastic noise process with defined characteristics, and a fixed spike threshold.

**Figure 4:**
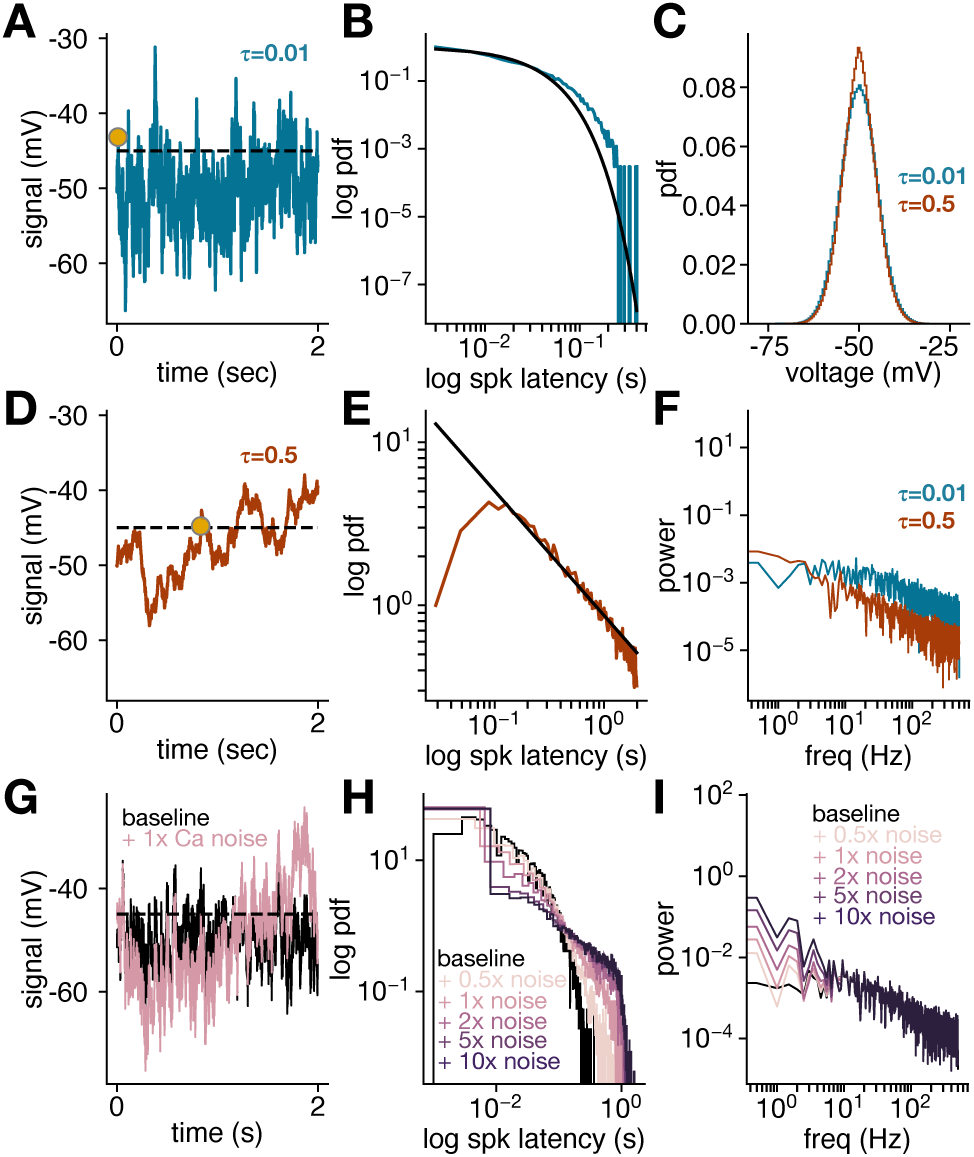
The statistics of membrane noise shapes stochastic spike timing in 5-HT neurons. **A**: Example simulation showing fast membrane potential noise represented by a fast (tau=0.01s) Ornstein-Uhlenbeck process. The dotted line represents spike threshold and the yellow dot represents the time of first-passage-crossing. **B**: Distribution of first-spike latencies across 20,000 independent initializations of OU noise. Black line represents an exponential fit. **C**: Distribution of simulated membrane voltages for fast and slow Ornstein-Uhlenbeck noise shown in **A** and **D**. **D**: Example simulation showing slow membrane potential noise represented by a slow (tau=0.5s) Ornstein-Uhlenbeck process. The dotted line represents spike threshold and the yellow dot represents the time of first-passage-crossing. **E**: Distribution of first-spike latencies across 20,000 independent initializations of OU noise. Black line represents a power-law fit. **F**: Spectral power of simulated membrane potential for fast and slow noise processes. **G**: Example simulations showing fast membrane potential noise (tau=0.01s Ornstein Uhlenbeck process, black) and fast membrane potential noise with calcium noise spectral power added in the frequency domain (pink). **H**: Distribution of first-spike latencies across 20,000 independent initializations of OU noise (black), and with increasing calcium noise spectral power added (pink). **I**: Spectral power of simulated membrane potential for fast noise process alone and with increasing calcium noise spectral power added.

Guided by our empirical findings showing that calcium channels altered the temporal structure of membrane noise, we first used *in silico* simulations to ask how the rate of noise fluctuations affected spike timing distributions. Ornstein-Uhlenbeck noise with a fast time constant (tau=0.01) generated rapidly fluctuating voltage signals that to reach spike initiation very quickly, with the spike times exponentially distributed and typically in the range of tens of milliseconds (Fig. 4A-B). Ornstein Uhlenbeck noise with a slow time constant (tau=0.5), and with a fixed membrane potential distribution compared to fast noise (Fig. 4C), generated time-varying signals that evolved more slowly and tended to show longer delays before reaching spike threshold. At the population level, these spike times were in the range of hundreds of milliseconds to seconds, and were power-law distributed (Fig. 4E). The spectral characteristics of these signals, revealed through a fast Fourier transform (Fig. 4F), showed that simulations with a slower noise time constant were specifically enriched in very low frequencies (0.5-2Hz). Together, these results provide a framework to understand how the distribution of spike initiation times depends on intracellular noise features. In particular, at subthreshold membrane potentials, noise with fast fluctuations tends to produce short spike latencies which are exponentially distributed over time, while noise with slower fluctuations produces more protracted spike latencies which are power-law distributed.

To provide a more direct comparison to the membrane noise observed in 5-HT neurons, we asked whether the observed spectral characteristics of this noise (Fig. 3M) could modulate spike latency distributions in a reliable manner. We thus first added the spectral components of the experimentally-observed calcium channel-dependent noise to the fast-fluctuating Ornstein-Uhlenbeck noise process presented in Fig. 4A. Consistent with its prominent low-frequency component, the addition of the spectral components of the calcium conductance-dependent noise generated slow fluctuations in the simulated signal (Fig. 4G, I). We determined how adding scaled components of the noise ranging from 0.5x to 10x (*i.e.,* weak to strong activation of calcium channels), affected spike initiation times. Adding the noise dramatically increased the average spike latency time, and prolonged the straight-line component of the spike latency distribution (in a log-log plot) to span two orders of magnitude, consistent with a power-law distribution of latencies (Fig. 4H). These results thus demonstrate that distinct spike latency distributions at the population level can be readily explained by stochastic noise processes occurring in single neurons, allowing these individual neurons to toggle between distinct latency ‘modes’ through activation or inactivation of ionic conductances. Ultimately, this suggests a mechanism by which populations of 5-HT neurons receiving overlapping synaptic inputs can effectively segregate these inputs through dynamic generation of synchronous (short-latency) or asynchronous (long-latency) spiking, utilizing the simple building blocks of stochastic noise processes and ionic conductances with defined spectral characteristics.

## Discussion

In this work, we demonstrate that a subpopulation of individual 5-HT neurons in DRN receives strong excitatory drive from both mPFC and LHb, and information from these two inputs can be multiplexed at the population level by cell-intrinsic regulation of spike timing. While synaptic inputs from LHb produce time-locked, low-latency spikes leading to population synchrony, mPFC inputs produce highly jittered, asynchronous spikes. We show that these two informational sources can be theoretically decoded by downstream circuits receiving sparse 5-HT afferents. Finally, we demonstrate that the delayed spiking pattern from mPFC inputs arise from a set of simple ionic mechanism: perithreshold calcium channels increase membrane noise and trigger stochastic, delayed spike threshold crossings. Together, these findings indicate that long-range inputs from mPFC and LHb can be multiplexed in local serotonergic circuits by cell-intrinsic regulation of spike synchrony.

### DRN 5-HT neurons multiplex information from multiple sources

Our results add to a body of work showing signals may be represented by precisely timed activity patterns in 5-HT neurons, including during complex behaviors. Paquelet et al. (2022) undertook miniscope calcium imaging of populations of 5-HT neurons during emotionally salient behaviors, finding highly correlated (synchronous) ensembles of neurons responding to sucrose or footshocks alongside non-selective ensembles. Li et al. (2016) demonstrated that 5-HT neurons encode reward signals, including a subpopulation that exhibits increased firing rate with millisecond precision. Cohen et al. (2015) reported that 5-HT neurons exhibited tonic firing rates that tracked block reward size, alongside phasic responses to direct rewards and punishments. These findings, taken alongside the results reported here, suggest a model where information about ongoing environmental stimuli, such as rewards or punishments, is encoded as synchronous population activity in DRN microcircuits, while higher-order task features or environmental statistics may be represented as uncorrelated firing of 5-HT neurons. Our dynamical model provides a putative explanation for both the uncorrelated spiking and synchrony shown to be present in populations of 5-HT neurons *in vivo* (Cohen et al., 2015; Paquelet et al., 2022). We expect that future work will help address the question of how synchronous and asynchronous activity evolves in larger populations of 5-HT neurons, and whether effective parsing of spiking in this manner provides meaningful information about task variables or internal variables.

### Calcium channel-dependent noise may provide a generalizable mechanism for separating inputs through differential spike timing

Our results demonstrate that 5-HT neurons exhibit calcium conductances that are activated in the perithreshold regime and add a broadband low-frequency component to membrane noise. How do synaptic inputs from mPFC preferentially activate these calcium conductances, while LHb inputs do not? Here we entertain two possibilities. First, a dendritic gradient of calcium channel expression, combined with localized targeting of calcium channel-rich dendritic segments by mPFC inputs, might produce mPFC-specific activation of calcium conductances. Indeed, differential targeting of dendritic segments by long-range inputs can profoundly shape spike output and integration in other brain areas (MacAskill et al., 2012), indicating they are a plausible substrate for the effects reported here. A second possibility is that mPFC synaptic inputs to 5-HT neuron dendrites could be more spatially clustered than LHb synaptic inputs, allowing mPFC inputs to exclusively trigger to dendritic calcium events when activated synchronously. Previous work mainly in cortex and hippocampus has shown that clustered synaptic activation generates several dendritic non-linearities, both in terms of calcium signaling and membrane depolarization (Lee et al., 2016; Higley & Sabatini, 2008; Mel, 1993; Losonczy & Magee, 2006). Future studies will be needed to disambiguate these and other possibilities.

The findings reported here add to previous studies that link the actions of specific ionic conductances to cellular and population spiking dynamics. For instance, electrosensory neurons in the weakly electrical fish exhibit Na^+^ channel noise, including “blips” that trigger action potentials (Marcoux et al., 2016). This noise was shown to boost slow excitatory synaptic inputs and the noise-triggered spikes enacted a stochastic process that prevented spike synchronization at a population level. The blips, somewhat analogous to the membrane noise observed here in DRN 5-HT neurons, were likely generated by clustered dendritic Na^+^ channels (Turner et al., 1994). We thus propose that intrinsic neuron noise driven by spontaneous opening of voltage-dependent Na^+^ or Ca^2+^ channels may be a common biophysical mechanism for decorrelating spiking outputs, even when the noisy cells are embedded in neuronal networks that perform different computations.

### Decoding of DRN input streams through differential 5-HT release in downstream brain areas

We used linear decoders to demonstrate that the observed spike timing patterns can theoretically be decoded by downstream circuits innervated by 5-HT axons. There are multiple biologically plausible mechanisms that could provide such a decoding capability. First, the short-term facilitation of 5-HT release (Lynn et al., 2022) could functionally segregate high-frequency and low-frequency spike trains, albeit from the same neuron, by differential 5-HT release dynamics. Such a short-term facilitation decoding mechanism has been shown to be theoretically viable in the electrosensory system of weakly electric fish (Middleton et al., 2011). Here, the viability of this decoding scheme would rely on LHb or PFC inputs changing the spiking statistics of single neurons at fine timescales corresponding to changes in input patterns. Recent work, showing that 5-HT neurons can emit action potentials transiently at high rates of ∼20-60Hz, appears to provide a plausible substrate for such a decoding mechanism (Li et al., 2016). A second biologically feasible mechanism involves a 5-HT affinity-based decoding scheme, determined by the differential 5-HT receptor subtype expression across cortical excitatory versus inhibitory neurons. In cortex, inhibitory 5-HT receptors of the 5-HT1A subtype are broadly expressed in pyramidal neurons, while excitatory 5-HT2A receptors are strongly expressed in apical dendrites of at least a subset of pyramidal neurons (Béïque et al., 2007; Jakab et al., 1999), while some neurons express both subtypes (Araneda & Andrade, 1991; Avesar & Gulledge, 2012). Moreover, fast-spiking and somatostatin interneurons can either express excitatory 5-HT2A or inhibitory 5-HT1A receptors, but typically not both (for review, see Puig et al., 2011). The differential affinity of 5-HT1A and 5-HT2A receptors for 5-HT (Peroutka et al., 1979; Nichols & Nichols, 2008), suggest a decoding scheme where 5-HT1AR-expressing neurons may be inhibited by both synchronous and asynchronous 5-HT axon release events, while 5-HT2AR-expressing interneurons and pyramidal neurons expressing 5-HT2ARs may be depolarized by only highly synchronous 5-HT release events due to lower receptor affinity. Such receptor binding rules in cortical microcircuitry could in principle sustain demultiplexing of information carried by 5-HT axon synchrony. Further work is needed to discriminate between these two hypothesized decoding schemes, and to establish the precise network mechanisms by which such decoding schemes would function.

Taken together, our results demonstrate that populations of 5-HT neurons receiving multiple sources of long-range synaptic input can effectively multiplex information, through predominantly cell-intrinsic mechanisms that regulate spike synchrony at the population level. These cell-intrinsic mechanisms involve stochastic membrane noise processes and differential calcium channel activation. More broadly, these results suggest that a high-dimensional population coding scheme such as synchrony-division multiplexing can plausibly arise from the autonomous, stochastic noise processes occurring in individual neurons (assuming synaptic volleys well separated in time), even in the absence of strong network-level interactions that are conventionally thought to shape information representations in cortex.

The effects of synchronous versus asynchronous activation of DRN 5-HT cells on individual pyramidal cells and interneurons will be amplified by local recurrent cortical circuitry. Further, DRN has widespread projections to cortex and the extensive reentrant cortical pathways will extend the effects of such inputs to cortical areas not directly associated with persistent cognitive states (PFC) or transient negative reward prediction errors (LHb). Extracting the signatures of the differential DRN 5-HT modulation of cortical activity from the baseline spiking activity generated by local and long-range cortical circuitry will be a major challenge for future studies.

## Acknowledgements

We thank all members of the Beique, Maler and Naud labs for discussion. This work was supported by grants from CIHR to Leonard Maler (221043) and Jean-Claude Beique (175319, 175325). Michael Lynn is grateful for graduate support provided by OGS and Queen Elizabeth II-GSST.

## Methods

### Animal use

SERTcre: B6.Cg-Tg(Slc6a4-cre)ET33Gsat/Mmcd

R26R-TdT: Rosa26R-TdTomato fluorescent protein (TdT) reporter mice

All experiments and procedures were performed in accordance with approved procedures and guidelines set forth by the University of Ottawa Animal Care and Veterinary Services. For this study, SERTcre mice were crossed with the R26R line to generate a SERTcre-TdT mouse line. Mice were kept on a 12:12 hour light/dark cycle, with access to food and water ad libitum unless otherwise noted. A mix of male and female SERTcre-TdT mice were used.

### Slice physiology

#### Stereotaxic Injections

All experiments and procedures were performed in accordance with approved procedures and guidelines set forth by the University of Ottawa Animal Care and Veterinary Services. For stereotaxic injections, mice were injected with 0.05 mg/kg buprenorphine and anesthetized by inhalation of isoflurane. Injections were performed using either a 10 μl Hamilton syringe with a 33 gauge needle, or a Nanoject machine with a glass micropipette. For all injections, 0.2 - 0.3 μl of virus was injected per coordinate, at a rate of 0.1 – 0.2 ul per minute. Stereotaxic co-ordinates are as follows: LHb (from Bregma skull landmark, in mm: AP −1.7, ML +/- 0.4, DV −2.7 to −3.0) and mPFC (from Bregma skull landmark, in mm: AP +2.5, ML +0.4, DV −1.5 to −1.8 from the surface of the brain).

For ChR2 (H134R) expression, 0.2-0.3 μl of AAV9.hSyn.hChR2(H134R)-mCherry.WPRE.SV40 (titre ∼1012 GC/mL) was injected bilaterally into either LHb or mPFC and animals were left to recover for 3 to 4 weeks prior to ex-vivo electrophysiological recordings.

For the dual opsin experiments (Chrimson/Chronos), 0.2-0.3 ul of AAV-Syn-ChrimsonR-TdTomato (Serotype:2/9) was injected bilaterally into LHb, and 0.2-0.3 ul of AAV-Syn-Chronos-GFP (Serotype:1) was injected bilaterally into mPFC. In some control experiments (marked in text and figures), only AAV-Syn-ChrimsonR-TdTomato (Serotype:2/9) was injected bilaterally into LHb to rule out spectral overlap between opsins. Animals were left to recover for 3 to 4 weeks prior to ex-vivo electrophysiological recordings.

### Slice preparation

*LED Photostimulation experiments (Fig. 1).* DRN containing brainstem slices were prepared from mice aged P40-P90. Acute slices were prepared similarly to previously described methods (Geddes et al., 2016). In brief, mice were anesthetized by isoflurane inhalation (Baxter Corporation, Canada) and sacrificed by decapitation. The brain was removed and brain slices were sectioned while immersed in ice-cold choline chloride-based cutting solution of the following composition (in mM): 119 choline-Cl, 2.5 KCl, 1 CaCl2, 4.3 MgSO4-7H2O, 1 NaH2PO4, 1.3 sodium L-ascorbate, 26.2 NaHCO3, and 11 glucose, and equilibrated with 95% O2, 5% CO2. Slices were recovered in a chamber containing standard Ringer’s solution of the following composition (in mM): 119 NaCl, 2.5 CaCl2, 1.3 MgSO4-7H2O, 1 NaH2PO4, 26.2 NaHCO3, and 11 glucose, at a temperature of 37°C, continuously bubbled with a mixture of 95% O2, 5% CO2. Slices were left to recover for 1 hour in a recovery chamber and equilibrate to a temperature of approximately 21°C until recordings were performed.

#### Calcium conductance and membrane noise experiments (*Fig. 3*)

DRN containing brainstem slices were prepared from mice aged P21-P40. To optimize the health of serotonin neurons for these sensitive experiments, acute slices were prepared using a modified NMDG two-bath cutting procedure, described in Ting et al. (2018). Briefly, mice were anesthetized by isoflurane inhalation (Baxter Corporation, Canada) and were transcardially perfused with ice-cold NMDG-based cutting solution with the following composition (in mM): 92 NMDG, 2.5 KCl, 1.2 NaH2PO4H, 30 NaHCO4, 20 HEPES, 10 MgSO4, 25 Glucose, 0.5 CaCl2, 5 sodium ascorbate, 2 thiourea, 3 sodium pyruvate, with pH adjusted to 7.3-7.4 with HCl and osmolality adjusted to 300-310 mOsm/kg with sucrose, and equilibrated with 95% O2, 5% CO2. Following perfusion, the brain was removed and 300um brain slices were sectioned in the same NMDG cutting solution. Brain slices were incubated for 0.5h in NMDG cutting solution at 37C, with NaCl concentration gradually increasing to 92 mM over the incubation time (details in Ting et al., 2018), and then transferred to a HEPES holding ACSF bath, at room temperature, with the following composition (in mM): 92 NaCl, 2.5 KCl, 1.2 NaH2PO4H, 30 NaHCO3, 2 MgSO4.7H2O, 2 CaCl2.2H2O, 20 HEPES, 25 Glucose, 5 Sodium ascorbate, 2 Thiourea, 3 Sodium pyruvate, equilibrated with 95% O2, 5% CO2. Slices were left to recover for 1 hour in a recovery chamber and equilibrate to a temperature of approximately 21°C until recordings were performed.

### Whole-Cell Electrophysiology

DRN neurons were visualized using an upright microscope: (1) Examiner D1 equipped with Dodt-contrast or differential-interference contrast (DIC) (Zeiss, Oberkochen, Germany; 40x/ 0.75NA objective) or, (2) BX51W1 equipped with DIC optics (Olympus, Center Valley, PA; 40x/ 0.80NA objective). 5-HT neurons were visually identified by TdTomato signal driven by the SERT promoter under EPI-fluorescence illumination (xCite Series 120pc). Whole-cell recordings were carried out using an Axon Multiclamp 700B amplifier, sampled at 10 kHz, digitized with an Axon Digidata 1440A (or 1550) digitizer and filtered at 2 kHz. Whole-cell recordings were performed using borosilicate glass patch electrodes (3-6 MΩ; World Precision Instruments, Florida) pulled on a Narishige PC-10 pipette puller (Narishige, Japan).

All LED photostimulation experiments (Fig. 1) were performed at room temperature (20C).

All calcium conductance and membrane dynamics experiments (Fig. 3) were conducted at a measured 30-32°C through the use of an in-line bath heater and stage heater. The purpose of these alterations were to replicate, as closely as possible, the membrane dynamics and channel kinetics that would be observed *in vivo*.

Slices received constant bath application of Ringer containing (in mM): 119 NaCl, 2.5 KCl, 1.3 MgSO4, 2.5 CaCl2, 1.0 NaH2PO4, 11 glucose, 26.2 NaHCO3, and 0.1 L-Tryptophan saturated with 95% O2 and 5% CO2 (pH = 7.3; 295-310 mOsm/L). An intracellular solution of the following composition was utilized: 115 mM potassium gluconate, 20 mM KCl, 10 mM sodium phosphocreatine, 10 mM HEPES, 4 mM ATP(Mg2+), and 0.5 mM GTP, pH 7.25 (adjusted with KOH; osmolarity, 280–290 mOsmol/L). For all voltage clamp recordings, access resistance was continuously monitored using a 125 ms, 2 mV hyperpolarizing pulse, at least 245 ms prior to stimulation. Recordings were discarded when the access resistance changed by >30%. Liquid junction potential was not compensated for. For current clamp experiments, during LED photostimulation, direct current injection was used to maintain membrane voltage at approximately ∼-50mV to allow a direct comparison between neurons in each stimulation condition (described in Fig. S2).

### LED Photostimulation

LED photostimulation was delivered using a PlexBright LED module and controller (465 nm; Plexon, Texas). Light was delivered through a 200 μm patch fiber cable with a bare fiber tip with ∼ 1 cm of glass exposed. For EPSC/EPSP stimulation, unless otherwise stated, optically evoked-PSCs were stimulated with single light pulses of 1-5 ms in length (individual trials 12- 15 seconds apart) and an irradiance of 792 mW/mm2.

### Drugs and Chemicals

Picrotoxin, CNQX and nickel (ii) chloride were purchased from Abcam (Cambridge, MA), Tocris Bioscience (Bristol, UK) and Sigma-Aldrich (St. Louis, MO), respectively. The concentration of drugs employed was as follows: Picrotoxin 100uM, CNQX 10uM, NiCl 250uM.

### Data Analysis

All electrophysiological recordings were analyzed offline in the Python computing environment using the Numpy and Scipy computational packages. All data are presented as means +/- SEM. n refers to the number of cells recorded unless otherwise stated. P values of less than 0.05 were considered statistically significant and are indicated with an asterisk(*).

### Applying linear classifiers to sub-threshold and spiking statistics

The approach to apply linear classifiers to the LHb and mPFC event data was as follows. We first describe methods to generate unbiased synthetic datasets of varying sample sizes from our physiological data, and second describe application of a linear classifier to this data.

#### Synthetic datasets

For a given number of samples and an event statistic (*e.g.* spike latency or EPSP amplitude), we generated 1000 synthetic datasets for both LHb and mPFC inputs. These synthetic datasets were generated by iteratively performing the following: 1) randomly sampling a neuron, then 2) randomly sampling a trial from that neuron and storing the event statistic from that trial. This was iteratively performed until the desired number of samples was reached. This sampling procedure ensured that each neuron was equally represented in the dataset. Next, the 1000 synthetic datasets were partitioned into either *training* or *test* datasets, with 5- fold cross-validation.

#### Linear classifier

We first performed standardization of the data using a min-max scaler, and then trained a support vector machine linear classifier on the *training* dataset using stochastic gradient descent learning (employing the implementation in scikit-learn). The classifier was then used to predict labels (LHb or mPFC) in the test dataset, with classification accuracy registered as the fraction of correct classifications across the entire test dataset. This linear classifier training and evaluation was then repeated for a total of 5 times (5-fold cross-validation) while rotating the held-out data in the test set. The classification accuracy was reported as the average accuracy across all 5 rotations of test and training data.

### Generation and injection of synthetic EPSCs (Fig. 3)

Synthetic EPSCs were generated as a waveform with instantaneous rise-time, and a monoexponential decay with *τ_decay_* that varied from 30ms to 500ms. These synthetic EPSCs were injected into the neuron in current clamp to elicit an evoked EPSP response. In Fig. 3C and Fig. S3, as various values of *τ_decay_* were presented to the same neuron, the total charge transfer to the neuron was held constant by scaling the amplitude of each EPSC such that the integral of current over time was calculated to be identical.

### Membrane noise analysis and spike threshold calculation (Fig. 3)

For the analysis of membrane potential dynamics, spikes were first identified using a fractional first-derivative method (Trinh et al., 2019; Azouz and Gray, 2000). Briefly, the maximum derivative of voltage over time, 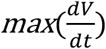, was calculated for each spike, and the spike threshold was identified as the first point in the depolarization where the voltage derivative reached 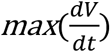. A spike comprised all points from spike threshold to 0.4s after the threshold; all such points were masked and not included in analysis of membrane voltage statistics or spectral characteristics. To calculate the spectral characteristics of membrane voltage, a 1-dimensional discrete fast Fourier transform of the voltage time series was computed, and the resulting spectrum was smoothed using a Savitzky-Golay filter with polynomial order 3.

### Stochastic membrane noise simulations (Fig. 4)

To investigate how distinct spike timing patterns could arise from stochastic fluctuations in membrane voltage, we simulated membrane voltage as a dynamical process and solved the level-crossing problem to infer how spike timing distributions depended on membrane dynamics. We considered a simplified scenario where membrane voltage (*v*_0_ = −50*mV*) was initialized close to spike threshold (*v_thresh_* = −4501). Membrane voltage evolved stochastically (described below) until spike threshold was reached. By repeating this over 20,000 simulations, we generated a distribution of level-crossing times (representing spike timing distributions), allowing us to assess how membrane potential dynamics were related to distinct regimes of spike timing.

We simulated the dynamics of membrane voltage as an Ornstein-Uhlenbeck noise with the following equation:

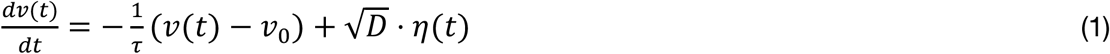

where *τ* is a constant denoting the correlation timescale of the noise (indicated in text, ranging from 0.01 to 0.5), *v*_0_ = −50 *mV* represents the resting membrane potential, and η(*t*) is a white noise term. 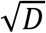 is a noise diffusion coefficient (scaling factor) that was varied across simulations with *τ* such that the quantity *σ*^2^ = *Dτ*/2 was constant (*σ*^2^ = 100); this produced fixed membrane potential distributions that were independent of *τ* as it was varied (Fig. 4B). The simulations proceeded until *v* reached the simulated spike threshold *v_thresh_* = −45*mV* at which point the time from simulation onset to spike threshold was recorded as a simulated spike latency.

Numerically, the Ornstein-Uhlenbeck noise process was computed using the following constant definitions and update rule, following the well-known algorithm derived by Gillespie (1996) for exact numerical simulation of the Ornstein-Uhlenbeck Process:

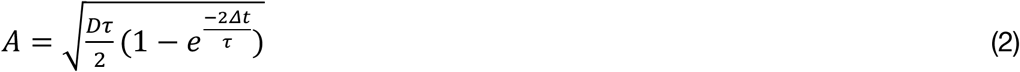

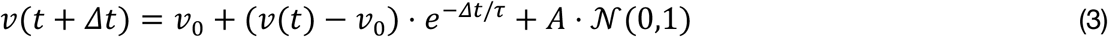

With a timestep of *Δt* = 0.001 *s*.

For simulations in which the spectral features of the noise were altered, we conducted simulations as described above, then performed a fast Fourier transform of the simulated voltage, altered the signal in the frequency domain, then transformed back into the time domain.

---

**Supplemental Figure 1:**
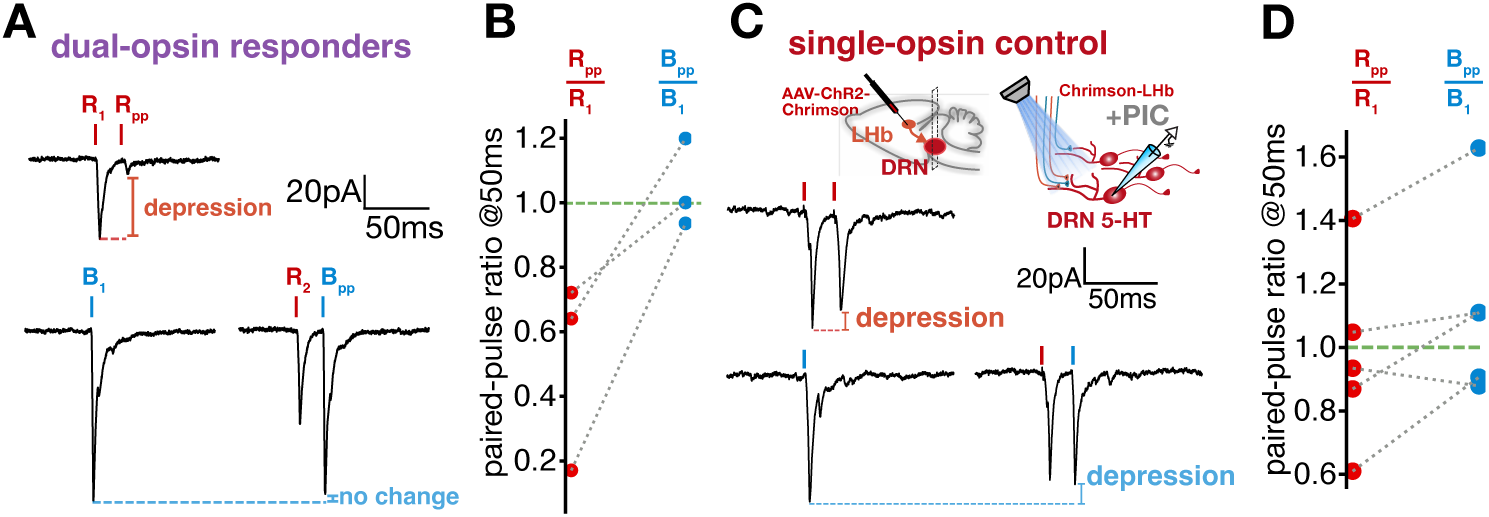
Dual-color opsin dissection of long-range inputs to 5-HT neurons. **A**: For 5-HT neurons that respond to both blue and red light, red paired-pulse stimuli produce depression (20Hz, top) while a red prepulse does not significantly affect the amplitude of a blue light-evoked current (bottom). **B**: Quantification of paired pulse ratio for red paired-pulse stimuli (red) and blue paired-pulse stimuli with red prepulse (blue). **C**: Single-opsin control with Chrimson injected in LHb. Red paired-pulse stimuli produce depression (20Hz, top) and a red prepulse significantly affect the amplitude of a blue light-evoked current (bottom). **D**: Quantification of paired pulse ratio for red paired-pulse stimuli (red) and blue paired-pulse stimuli with red prepulse (blue).

**Supplemental Figure 2:**
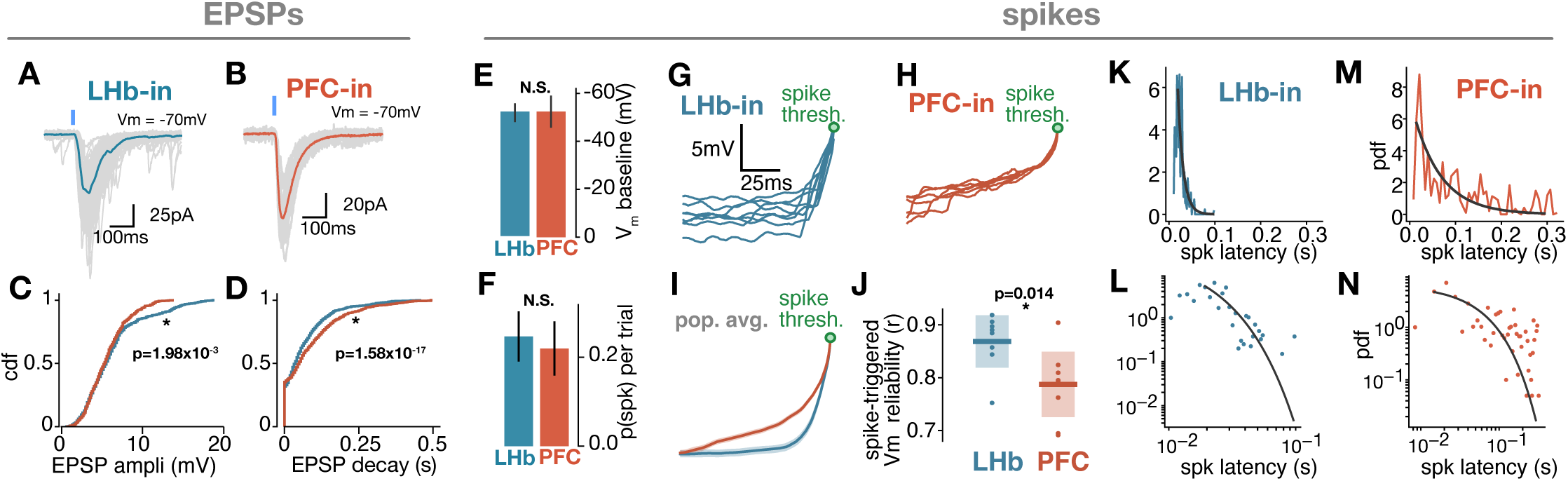
Quantifying features of EPSPs and spikes elicited by LHb and PFC inputs to 5-HT neurons. **A-B**: Example traces in voltage clamp of light-evoked EPSCs in 5-HT neurons. **C**: Cumulative distribution of EPSP amplitudes of all LHb-evoked events (blue) and PFC-evoked events (red) (K-S test, n=631 events from 9 neurons, LHb; n=619 events from 9 neurons, PFC). **D**: Cumulative distribution of EPSP decay of all LHb-evoked events (blue) and PFC-evoked events (red) (K-S test, n=631 events from 9 neurons, LHb; n=619 events from 9 neurons, PFC). **E**: Mean baseline membrane potential before evoked spikes (n=9 neurons for LHb and PFC groups, bars indicate S.E.M; p>0.05, t-test). **F**: Mean probability for photostimulation to trigger a spike (n=9 neurons for LHb and PFC groups; bars indicate S.E.M; p>0.05, t-test). **G-H:** Trial-by-trial voltage trajectories to spike threshold in two example neurons receiving suprathreshold long-range LHb (**G)** or PFC (**H**) input. **I**: Population average of voltage trajectories to spike threshold (n=194 spikes from 9 neurons, LHb; n=203 spikes from 9 neurons, PFC; shaded lines denote S.E.M). **J:** Reliability of voltage trajectory to spike threshold across trials, for LHb and PFC inputs (Reliability is computed as pairwise pearson r coefficient of voltage trajectory across all trials within a neuron; n=9 neurons for PFC and LHb, p=0.014, t-test). **K-N**: Probability distribution of spike times triggered by PFC (red) and LHb (blue), shown with regular scaling (**K-M**) and log-log scaling (**L-N**). Black lines indicate exponential fit.

**Supplemental Figure 3:**
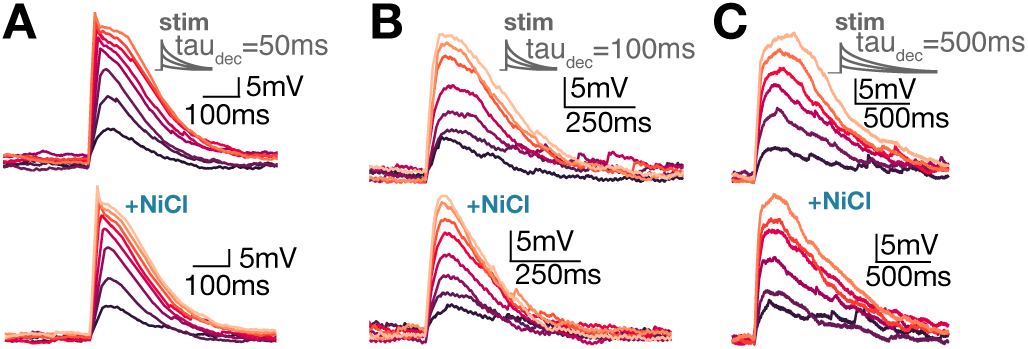
Low-threshold calcium channels modulate the kinetics of EPSPs in 5- HT neurons. **A**: Example traces in current-clamp showing that injection of fast (tau=50ms) synthetic EPSCs produce EPSP responses that are modulated by bath application of NiCl. **B**: Example traces in current-clamp showing that injection of fast (tau=100ms) synthetic EPSCs produce EPSP responses that are modulated by bath application of NiCl. **C**: Example traces in current-clamp showing that injection of fast (tau=500ms) synthetic EPSCs produce EPSP responses that are modulated by bath application of NiCl.

**Supplemental Figure 4:**
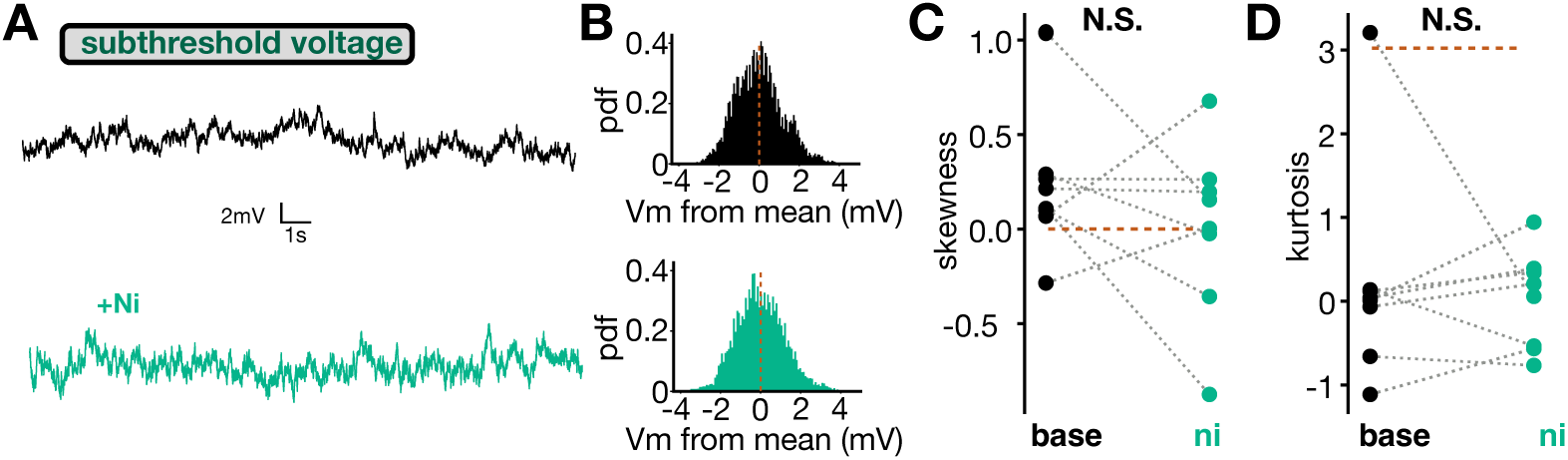
Subthreshold noise properties are not affected by calcium channels in 5-HT neurons. **A**: Example traces of a single neuron recorded in current-clamp during injection of constant subthreshold depolarizing current, before (top) and after (bottom) application of NiCl. ‘Subthreshold’ is defined as traces recorded during a current step just below spike threshold. **B**: Probability distribution of membrane potentials for an example neuron before (top) and after (bottom) application of NiCl. **C**: Membrane potential distribution skewness is not significantly affected by bath application of NiCl (n=7 neurons, paired T-test). **D**: Membrane potential distribution kurtosis not is significantly affected by bath application of NiCl (n=8 neurons, paired T-test).

**Supplemental Figure 5:**
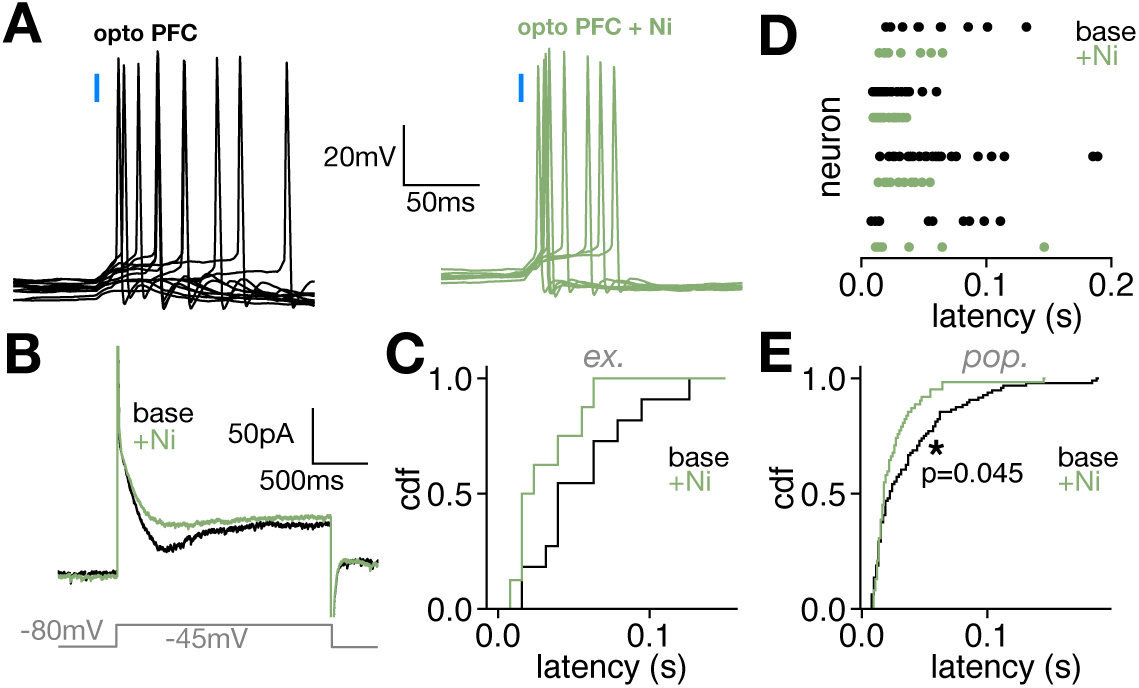
Low-threshold calcium channels contribute to spike timing properties of long-range PFC input to 5-HT neurons. Panels **A-C** depict results from a single representative 5-HT neuron. **A**: Representative traces from 5-HT neuron in current clamp depicting high-latency spiking in response to optogenetic PFC axon stimulation (left), and lower latency spiking after bath application of NiCl (right). **B**: Representative averaged trace in voltage clamp depicting inward current in response to a −45mV depolarizing step (black), which is abolished by bath application of NiCl (green). **C**: Cumulative distribution of spike latency before (black) and after (green) NiCl application from a single 5-HT neuron. **D**: Timing of spikes triggered by photostimulation of long-range PFC axons before (black) and after (green) bath application of NiCl. Each dot is a spike and each row depicts one neuron (n=4 neurons). **E**: Cumulative distribution of spike times triggered by photostimulation of PFC axons before (black) and after (green) bath application of NiCl (Kolmogorov-Smirnov test, p=0.045, n=96 spikes (base) and 62 spikes (Ni) from 4 neurons).

